# Response of *rhizobial* strains on biochemical traits and nutrient uptake in Mungbean Vigna radiata L. Wilczek under moisture stress

**DOI:** 10.1101/2021.03.16.435312

**Authors:** Sapna, K. D Sharma

## Abstract

The present study was conducted to assess the biochemical responses and nutrient uptake in response to *rhizobial* inoculations in mungbean, and to screen the *rhizobial* isolates for drought tolerance. A field experiment was designed in randomized block design and replicated thrice during *kharif* 2016 at Crop Physiology Field Area, CCS, Hisar. The experiment consisted of two levels of treatments (1) without inoculation (only RDF) and (2) with inoculation (RDF with combination of five *rhizobial* strains viz. *Vigna* 703 + PSB strain P-36, MR 63, MR 54, MB 17a and MH 8b2) and two environments i.e. rainfed (no post sowing irrigation) and irrigated. Membrane stability index, leghaemoglobin content, chlorophyll content reduced by 17.7 %, 24.5% and 2.9% resp. under rainfed conditions while the plants inoculated with *rhizobial* isolate MR63 and MB 17a showed greater chlorophyll content (20.2% and 16.2%), LHb (29.1% and 22.9%) and MSI (19.4% and 17.9%) and enhanced nutrient uptake over RDF.

## Introduction

Mungbean is one of the 3^rd^ most important legume after chickpea and pigeon pea. It is a vital source of protein, carbohydrates, minerals, fibers, antioxidants like flavonoids (Quercetin-3-Oglucoside) and phenolics for vegetarian in dietary habit (Guo *et al*., 2012). It is a short-duration legume (<65 days), that makes it an ideal crop for catch cropping, relay cropping and intercropping. Despite being an economically important crop, overall production of mungbean is low due to abiotic and biotic stresses (Bangar *et al*., 2018). Among the various environmental stresses, the major factor limiting the crop yield is the amount of moisture available to the crop during the growing season. In India, about 68% of net sown area (140 million hectares) is reported to be vulnerable to drought conditions. Mungbean when cultivated in post rainy season faces water stress at various stages of crop growth (Rambabu *et al*., 2016). The effects of water stress in mungbean has been found to be more pronounced on reproductive stage than other growth stages and yield is drastically reduced (Majeed *et al*. 2016).

Water stress affects various physiological processes associated with growth, development, and economic yield of a crop (Reboucas *et al*., 2017). Water deficit alters normal uptake of different essential nutrients that causes reduced plant yield. Cell membranes are one of the first sites of damage and it is generally accepted that the maintenance of their integrity and stability under water stress conditions is a major indicator of drought tolerance in plants. The occurrence of stress indicated by cell membrane injuries leads to an increased leakage of electrolytes (Tint *et. al*., 2011). Chlorophyll is the main chloroplast component for photosynthesis and substantial chlorophyll content has a constructive association with photosynthetic rate (Shobhkhizi *et al*. 2014). A study by Mafakheri *et. al*. (2010) indicated that, water deficit results in negative impact in plants as majority of chlorophyll are lost.

*Rhizobia* influence the physiological growth conditions of mungbean by nitrogen-fixing symbiosis and thus increasing its chlorophyll contents (Anjum *et. al*. 2006). However, N2 deficiency give a negative response in plants by showing symptoms of yellowing which demonstrate chlorophyll deterioration has occurred in plants. The bacteria lodging around the plant roots (*rhizobacteria*) are also more versatile in transforming, solubilizing and mobilizing other essential nutrients compared to those from bulk soils (Hayat *et al*., 2010). Therefore, the *rhizobacteria* are the dominant deriving forces in recycling the soil nutrients and consequently, they are crucial for soil fertility (Glick, 2012). Keeping in view numerous manifestations of a beneficial action of *Rhizobium* bacteria on plants, present investigation screens the various *rhizobial* strains in terms of biochemical traits and nutrient uptake in Mungbean (Vigna radiata L. Wilczek) under moisture stress.

## Materials and Methods

A field experiment was conducted during the rainy (*kharif*) season of 2016 in the drought plots at Crop Physiology Field Area, Department of Agronomy, CCS Haryana Agricultural University, Hisar (Haryana) in western side at 29^0^10’ North latitude, 75^0^46’ East longitude and at an altitude of 215.2 metres above mean sea level. The plot size for each treatment was 2.5 x 1.8 m (six rows of 2.5 m length with 30 cm spacing) with rainout shelters facilities. The soils had alkaline pH (8.60), available Nitrogen 112.7 kg/ha and available Phosphorus 12.0 kg/ha. Mungbean *rhizobial* isolates were procured from Department of Microbiology, CCS Haryana Agricultural University, Hisar and the seed inoculation was done 2-3 h before sowing. The experiment included 6 treatment combinations (RDF and rhizobial strains) i.e. Recommended dose of fertilizer (RDF), RDF + *Rhizobium sp.(vigna)703* + PSB strain P-36, RDF + MR 63, RDF + MR 54, RDF + MB 17a and RDF + MH 8b2. The variety MH-421 of mungbean was used as the test crop and the sowing was done on 27^th^ June, 2016. Recommended dose of fertilizers (20 kg N and 40 kg P_2_O_5_) and crop protection measures was adopted as per package and practices.

Leaf discs (0.05 g) were washed, blotted dry and dipped in test tubes containing 5 ml of dimethyl sulfoxide (DMSO) overnight as described by Sawhney and Singh (2002). The extracted chlorophyll in DMSO was estimated by recording its absorbance at 663 and 645 nm, respectively

The assessment of membrane stability was done by the procedure of Dionisio-Sese and Tobita (1998). 100 mg of leaf tissue was taken separately in 20 ml test tubes containing 10 ml of de-ionized water and kept in 20 ml vials containing 10 ml de-ionized water at 25°C. After 4 h, the electrical conductivity (EC) of the solution was measured and designated as EC_a_. Then the samples were kept in boiling water bath for 1 h to achieve total killing of the tissue. After cooling, the EC of the solution was again measured and designated as EC_b_.

Leghaemoglobin (LHb) was determined by the method of Hartree (1995) with some modifications. This method was based on the conversion of haematin in pyridine haemochromogen. The whole plant nodules of known weight were homogenized in 5.0 ml of chilled 0.1 M phosphate buffer (pH7.0) in pestle mortar placed in ice bath. The extract was centrifuged at 5000 ppm for 30 minutes. At 4°C on refrigerated centrifuge 5.0 ml of supernatant was used for leghaemoglobin estimation.

The content of nitrogen and phosphorus were estimated in the seeds and stover of plant after harvesting. Digestion of plant material was done by adding di-acid mixture (H_2_SO_4_: HClO_4_ in 9:1 ratio). The nitrogen content was determined by using the method as Lindner (1994) and the phosphorus content was determined by vando-molybdophosphoric acid yellow colour method as described by Koenig and Johnson (1942).

## Results and Discussion

Results presented in Table 1 showed the total chlorophyll content decreased (2.9%) under water stress over irrigated. Such decrease in chlorophyll content in the leaves of plants may be attributed to the high rate of chlorophyll degradation more than its biosynthesis under water stress conditions (Raina *et al*. 2016). Application of *rhizobial* isolates increased the total chlorophyll content and the increase was more in RDF + MR 63 (4.91 mg/g/FW) followed by RDF + MB 17a (4.70 mg/g/FW) over RDF (3.91 mg/g/FW). The increased chlorophyll in *rhizobial* inoculated treatments may be due to improved plant water status resulted in reduced chlorophyll degradation and enhanced chlorophyll formation. Similar effect of *rhizobial* treatments on chlorophyll content was observed by Tairo *et al*. (2017)

**Table 1:** Effect of soil moisture and rhizobial isolates on biochemical parameters of mungbean

| Treatment | Chlorophyll content |  |  | MSI (%) |  |  | LHB content |  |  |
| --- | --- | --- | --- | --- | --- | --- | --- | --- | --- |
|  | IR | RF | Mean | IR | RF | Mean | IR | RF | Mean |
| **RDF (N and P)** | 3.93 | 3.88 | 3.91 | 13.04 | 10.82 | 11.93 | 1.92 | 1.40 | 1.66 |
| **RDF + Vigna703+P-36** | 4.22 | 4.14 | 4.18 | 15.81 | 12.16 | 13.99 | 2.03 | 1.46 | 1.75 |
| **RDF + MR 63** | 5.00 | 4.82 | 4.91 | 20.91 | 17.84 | 19.38 | 2.65 | 2.03 | 2.34 |
| **RDF + MR 54** | 4.31 | 4.17 | 4.24 | 17.32 | 14.35 | 15.84 | 2.23 | 1.68 | 1.96 |
| **RDF + MB 17a** | 4.75 | 4.64 | 4.70 | 19.25 | 16.57 | 17.91 | 2.42 | 1.89 | 2.16 |
| **RDF + MH 8b2** | 4.34 | 4.14 | 4.24 | 17.93 | 14.72 | 16.33 | 2.13 | 1.64 | 1.89 |
| **Mean** | 4.43 | 4.30 |  | 17.38 | 14.41 |  | 2.23 | 1.68 |  |
| **CD (P=0.05)** | E= 0.10 ; T=0.22;<br>E×T=0.37 |  |  | E= 1.19; T=2.04;<br>E×T= 2.71 |  |  | E= 0.15; T= 0.35;<br>E×T=0.48 |  |  |

\* E=Environment, T= Treatment, NS=Non significant, IR= Irrigated, RF= Rainfed

**Table 2:** Effect of soil moisture and rhizobial isolates on nitrogen and phosphorus content in straw and seeds of mungbean

| Treatment | N(%) in Seed |  |  | N(%) in Straw |  |  | P(%) in Seed |  |  |
| --- | --- | --- | --- | --- | --- | --- | --- | --- | --- |
|  | IR | RF | Mean | IR | RF | Mean | IR | RF | Mean |
| **RDF (N and P)** | 3.44 | 3.30 | 3.37 | 1.84 | 0.98 | 1.41 | 0.29 | 0.19 | 0.24 |
| **RDF + Vigna703+P-36** | 3.56 | 3.46 | 3.51 | 1.96 | 1.07 | 1.52 | 0.34 | 0.20 | 0.27 |
| **RDF + MR 63** | 4.06 | 3.86 | 3.96 | 2.40 | 1.45 | 1.93 | 0.55 | 0.30 | 0.43 |
| **RDF + MR 54** | 3.62 | 3.48 | 3.55 | 2.18 | 1.18 | 1.68 | 0.38 | 0.22 | 0.30 |
| **RDF + MB 17a** | 3.82 | 3.64 | 3.73 | 2.27 | 1.39 | 1.83 | 0.46 | 0.26 | 0.36 |
| **RDF + MH 8b2** | 3.61 | 3.47 | 3.54 | 2.13 | 1.22 | 1.68 | 0.38 | 0.24 | 0.31 |
| **Mean** | 3.69 | 3.54 |  | 2.13 | 1.22 |  | 0.40 | 0.24 | 0.32 |
| **CD (P=0.05)** | E=0.06;T=0.01;<br>E×T=0.07 |  |  | E=0.03; T=0.06;<br>E×T=0.08 |  |  | E=0.01; T=0.01;<br>E×T=0.02 |  |  |

The results revealed that membrane stability index (MSI) significantly decreased during water stress (17.38 to 14.7%) as presented in table-1 might be due to reduced cell water potential. Our result is in agreement with the findings of previous researchers who suggested that membrane integrity and stability are two important factors contributing to drought tolerance capacity and yield stability of crops (Diego *et al*. 2012). Among the *rhizobial* isolates, RDF + MR 63 showed maximum membrane stability (19.4 %) followed by RDF + MB 17a (17.9%), while, the minimum MSI was observed in plants treated with RDF (11.9 %) accompanied by RDF + Vigna703+ P-36 (14.0%) irrespective of the environment. The results indicated that inoculation with *rhizobium* spp. gives tolerance to plants under drought stress, because the permeability of inoculated crops decreased under drought stress. The lowest reduction of MSI in *rhizobial* strain MR 63 and MB 17a revealed its higher capacity to tolerate the adverse effects of drought. The similar effect of *rhizobium* inoculation on *P. vulgaris* under drought was also reported by Quinto *et al*. (2015).

Highest leghaemoglobin content in root nodules was recorded in plants grown under irrigated condition (2.23 mg/g DW) as compared to rainfed (1.68 mg/g DW) at the flowering stage (Table 1). Water stress induced onset of nodule senescence may probably be responsible for decline in leghaemoglobin content and consequently inhibition of N2-fixation (Kumar and Kuhad, 2003). Among the treatments, the highest value of leghaemoglobin content was observed in *rhizobial* strain MR 63 (2.34 mg/g DW) followed by MB 17a (2.16 mg/g DW). The leghaemoglobin content in RDF was lowest (1.66 mg/g DW) and was at par with Vigna 703+P 36 (1.75 mg/g DW) and RDF + MH 8b2 (1.89 mg/g DW). Our results are in accordance with the findings of Rodrigues *et al*. (2013) who reported the similar increase in LHb in response to *rhizobial* inoculations.

### Nitrogen and phosphorus content

Water stress significantly decreased the nitrogen and phosphorus content in both seed and straw over normal irrigation (Table 1). Decrease in N and P content under moisture stress from 3.69% to 3.54% & 0.40% to 0.24% and 2.13% to1.22% & 0.15% to 0.11% in seed and straw resp. over the irrigated control, irrespective of the *rhizobial* treatments.. Nutrients from the soil reached to the surface of root by mass flow and diffusion processes. Mass flow and diffusion processes are positively correlated with moisture content of the soil. Movement of nutrients thought the plant is also associated with soil water content (Khaton *et al*., 2016). Thus, the greater content and uptake of N and P in seed and straw was observed under irrigated environment as compared to rainfed. Our results are supported by the findings of Ali *et al*. (2016) and Mohammad *et al*. (2017). Maximum nitrogen and phosphorus content was 3.96 & 0.43% in seed and 1.93 & 0.16 % in straw observed in plants treated with *rhizobial* isolate MR 63.

## Conclusion

In the present study, the five *rhizobial* isolates namely V*igna* 703 + PSB strain P-36, MR 63, MR 54, MB 17a and MH 8b2 were evaluated for drought tolerance using biochemical traits and nutrient uptake in seed and straw. Based on the findings of this experiment, plants inoculated with *rhizobial* isolate “MR63 and MB 17a”shows greater chlorophyll content (20.2% and 16.2%), LHb (29.1% and 22.9%) and MSI (19.4% and 17.9%) and enhanced nutrient uptake over RDF

